# Optimising Spectronaut search parameters to improve data quality with minimal proteome coverage reductions in DIA analyses of heterogeneous samples

**DOI:** 10.1101/2023.10.11.561927

**Authors:** Christa P. Baker, Roland Bruderer, James Abbott, J. Simon C. Arthur, Alejandro J. Brenes

## Abstract

Data independent acquisition has seen breakthroughs that enable comprehensive proteome profiling using short gradients. As the proteome coverage continues to increase, the quality of the data generated becomes much more relevant. Using Spectronaut, we show that the default search parameters can be easily optimised to minimise the occurrence of false positives across different samples. Using an immunological infection model system to demonstrate the impact of adjusting search settings we analysed mouse macrophages and compared their proteome to macrophages spiked with *Candida albicans*. This experimental system enabled the identification of ‘false positives’ since *Candida albicans* peptides and proteins should not be present in the mouse only samples. We show that adjusting the search parameters reduced ‘false positive’ identifications by 89% at the peptide and protein level, thereby considerably increasing the quality of the data. We also show that these optimised parameters incur a moderate cost, only reducing the overall number of ‘true positive’ identifications across each biological replicate by <6.7% at both the peptide and protein level. We believe the value of our updated search parameters extends beyond a two-organism analysis and would be of great value to any DIA experiment analysing heterogenous populations of cell types or tissues.

## Introduction

Data independent acquisition (DIA) has become a popular method to analyse label free proteomes at scale and without a considerable compromise in depth (1, 2). Unlike Data Dependent Acquisition (DDA), DIA does not select a subset of precursor ions to be fragmented for MS2 analysis, instead it fragments all ions over a specific M/Z window (3–5). Consequently, the spectra derived from DIA data are more complex and convoluted than those derived from DDA. This initially meant that that software tools (6–8) required libraries to be generated, either spectra or peptide centric, in order to analyse and match DIA data. Library generation became a barrier, due to cost and time constraints, hence, the development of library free DIA was a major advance that helped to popularise the method. Today, library free single-shot DIA workflows are a standard technique for large-scale comprehensive proteomics analyses.

DIA can now achieve near comprehensive proteome coverage while maintaining scalability (1, 2, 9), as such DIA-based analyses to study the immune system have also become more common and important (10–18). In this type of study, it is commonly necessary to analyse a host organism, frequently *Mus musculus* (Mouse) which are good models to study immunological networks which are broadly applicable to humans, and an invading pathogen such as *Candida albicans (C. albicans)* This type of analysis introduces additional complications to the standard proteomic workflows. Analysing biological samples that include more than one species increase the search space and introduce peptide mapping issues when there is sequence homology between the different peptidomes (19). To address this challenge, we explored the impact of different Spectronaut settings on the identification of false positives in samples with lysates derived from 1 or from 2 species. Here we analysed the proteomes of Mouse bone marrow derived macrophages (BMDM) in the presence or absence of the pathogen *C. albicans*.

Macrophages are a key cell of the innate immune system, which are known to help control infections and disease, such as *C. albicans*(20–23)*. C. albicans* is an opportunistic pathogen found in most healthy adults, but can quickly turn into difficult to treat, systemic infection if the immune system is suppressed, and carries a 46-75% mortality rate (21–23). Macrophages are one of the first immune cells to respond to and kill *C. albicans* via pathogen recognition receptors, resulting in phagocytosis of *C. albicans* yeast as well as macrophage activation resulting in the production of reactive oxygen species to aid killing and inflammatory cytokines to attract other immune cells to the site of infection (24–31).

Our results show that despite limited homology between the two species, a library free workflow using directDIA with the default search settings incorrectly identified over 1,000 *C. albicans* peptides in the non-spiked BMDM samples. Within this study we share optimised search parameters for Spectronaut that minimise the number of misidentifications, i.e. *C. albicans* peptides detected in the Mouse only samples, at a reduced penalty to overall proteome coverage. We believe our suggested setting will provide alternatives for more high-stringency, robust identification results for library-free DIA for multi-species that is also applicable to the analysis of heterogenous populations, be it different cell types or tissues.

## Experimental section

### Animal and Cell Culture

All Mouse work was done at the University of Dundee, following the UK Home Office licence (PAAE38C7B) and approved by the Ethical Review and Welfare Committee at the University of Dundee. C57Bl6/J mice were obtained from Charles River Laboratories, UK. The animals were maintained in accordance UK and EU regulations and were kept in specific pathogen free environment in individually ventilated cages. Nonbreeding mice were maintained in same sex cage groups, with access to water and food (R&M1 SDS, Special Diet Services). Animals were kept in light and dark (12/12-hours) cycle rooms at 21°C with 45-65% humidity. Mice were culled using in cage increased CO_2_ followed by confirmation of death by cervical dislocation.

### Murine bone marrow derived macrophage cell culture

Bone marrow derived macrophages (BMDMs) were cultured in L929 media as detailed. On day 0, Bone Marrow was flushed with PBS from the femurs and tibias of a mouse using a 25G needle and 20 mL syringe, onto a 90 mm bacteriological grade plate (Thermo Scientific, 101R20). Once flushed, the bone marrow was then filtered through a 100 μm strainer, into a 50 mL falcon tube, and centrifuged at 400 *g* for 5 minutes. The bone marrow was then resuspended in 100mL of L929 conditioned media (DMEM (FISHER Invitrogen, Cat. 11510416), supplemented with 10% heat-inactivated FBS (Labtech), 10% week 1 L929 conditioned media, 10% week 2 L929 conditioned media, 500 mL 100 μg/mL Penicillin and 100 μg/mL Streptomycin (GiBCO Life Technologies), 1% sodium pyruvate (Lonza), 1% 1xnonessential amino acids (Lonza), 50 μM 2-mercaptoethanol, 1% GlutaMAX (GIBCO Life Technologies) and 1% HEPES (Lonza)). Week 1 and Week 2 L929 media was collected from L929 fibroblasts in the media recipe listed above; however, without the L929 media supplements. Cells were cultured for 7 days on bacterial grade plates (Thermo Scientific, 101R20) at 37°C and 5% CO_2_. After 7 days, BMDM were detached by scraping in PBS with 1% EDTA, counted using a hemocytometer and replated at 1×10^6^ cells in 2 mL of L929 media on a 6-well TC treated plate (Geiner, 657-160), and allowed to seed overnight. The following morning, cells were gently washed three-times with PBS, and then cells were ready for lysis for proteomics.

### Candida albicans Cell Culture

Frozen SC5314 *C. albicans* strain (20% glycine, 80% YPD media containing *C. albicans* yeast) was used to streak a YPD agar plate (YPD medium + 2% agar) in sterile conditions and incubated overnight at 30°C. After distinct colony growth, a single colony was picked using pipette tip and cultured in 5 mL YPD media broth overnight in 30°C at 200 rpm.

### Lysis for proteomics

For this experiment, 3 mice were used, and one colony cultured from *C. albicans*. BMDMs and *C. albicans* were lysed separately using 400 μL/1×10^6^ cells with 5% SDS (20% SDS Sigma, 05030), 10mm TCEP (0.5 M TCEP ThermoFisher Scientific, 77720), 50 mM TEAB (1M TEAB ThermoFisher Scientific, 90114) and HiPerSolv Water for HPLC (VWR, 83650.320). Lysis and proteomic sample preparation are as described (17). Briefly, after lysis, lysates were boiled at 100°C for 5 minutes, and then sonicated before protein concentration was calculated using the EZQ protein quantification kit (Thermo Fisher Scientific, R33200). Each of the 3 Mouse biological replicates were separated into two aliquots each containing 200 μg of protein. One aliquot contained only murine BMDMs (non-spiked) while the other aliquot contained BMDMs spiked with 50 μg of *C. albicans* protein (spiked). Tryptic peptides were generated by the S-Trap Method using S-Strap: Rapid Universal MS Sample Prep (Co2-mini,Protifi) and Trypsin Gold (Promega, V5280). Samples were then vacuum dried and resuspended in 50 μL of 1% formic acid (Thermo Fisher Scientific, 695076). Sample peptides were calculated via CBQCA quantification kit (Thermo Fisher) and were then ready for MS-analysis.

### Liquid Chromatography-Mass Spectrometry

Samples were analysed on an Orbitrap Exploris 480 (ThermoFisher), in DIA mode (17). For each sample, 1.5 *µg* of peptide was analysed on the Exploris 480 coupled with a Dionex Ultimate 3000 RS (Thermo Scientific). LC buffers prepared as follows: buffer A (0.1% formic acid in Milli-Q water (v/v)) and buffer B (80% acetonitrile and 0.1% formic acid in Milli-Q water (v/v)). 1.5 μg aliquot of each sample were loaded at 15 μL/minute onto a trap column (100 μm × 2 cm, PepMap nanoViper C18 column, 5 μm, 100 Å, Thermo Scientific) equilibrated in 0.1% trifluoroacetic acid (TFA). The trap column was washed for 5 min at the same flow rate with 0.1% TFA then switched in-line with a Thermo Scientific, resolving C18 column (75 μm × 50 cm, PepMap RSLC C18 column, 2 μm, 100 Å). The peptides were eluted from the column at a constant flow rate of 300 nl/min with a linear gradient from 3% buffer B to 6% buffer B in 5 min, then from 6% buffer B to 35% buffer B in 115 min, and finally to 80% buffer B within 7 min. The column was then washed with 80% buffer B for 6 min and re-equilibrated in 3% buffer B for 15 min. Two blanks were run between each sample to reduce carry-over. The column was kept at a constant temperature of 50°C at all times.

The data was acquired using an easy spray source operated in positive mode with spray voltage at 1.9 kV, the capillary temperature at 250°C and the funnel RF at 60°C. The MS was operated in data-independent acquisition (DIA) mode. A scan cycle comprised a full MS scan (m/z range from 350–1650, with a maximum ion injection time of 20 MS, a resolution of 120,000 and automatic gain control (AGC) value of 5 × 10 ^6^). MS survey scan was followed by MS/MS DIA scan events using the following parameters: default charge state of 3, resolution 30.000, maximum ion injection time 55 MS, AGC 3 × 10 ^6^, stepped normalized collision energy 25.5, 27 and 30, fixed first mass 200 m/z. The inclusion list (DIA windows) and windows widths are shown in Table 1. Data for both MS and MS/MS scans were acquired in profile mode. Mass accuracy was checked before the start of samples analysis.

### DIA isolation windows

**Table 1:** DIA windows.

| Window | Window start (m/z) | Window width | Window overlap | Window | m/z | Isolation window | Window Overlap |
| --- | --- | --- | --- | --- | --- | --- | --- |
| 1 | 349.975 | 66.8 | 0.525 | 24 | 663.25 | 14.5 | 0.5 |
| 2 | 416.25 | 13.5 | 0.5 | 25 | 677.25 | 13.5 | 0.5 |
| 3 | 429.25 | 11.5 | 0.5 | 26 | 690.25 | 13.5 | 0.5 |
| 4 | 440.25 | 12.5 | 0.5 | 27 | 703.25 | 14.5 | 0.5 |
| 5 | 452.25 | 11.5 | 0.5 | 28 | 717.25 | 16.5 | 0.5 |
| 6 | 463.25 | 11.5 | 0.5 | 29 | 733.25 | 15.5 | 0.5 |
| 7 | 474.25 | 11.5 | 0.5 | 30 | 748.25 | 16.5 | 0.5 |
| 8 | 485.25 | 10.5 | 0.5 | 31 | 764.25 | 18.5 | 0.5 |
| 9 | 495.25 | 11.5 | 0.5 | 32 | 782.25 | 17.5 | 0.5 |
| 10 | 506.25 | 11.5 | 0.5 | 33 | 799.25 | 18.5 | 0.5 |
| 11 | 517.25 | 11.5 | 0.5 | 34 | 817.25 | 19.5 | 0.5 |
| 12 | 528.25 | 10.5 | 0.5 | 35 | 836.25 | 20.5 | 0.5 |
| 13 | 538.25 | 11.5 | 0.5 | 36 | 856.25 | 20.5 | 0.5 |
| 14 | 549.25 | 10.5 | 0.5 | 37 | 876.25 | 22.5 | 0.5 |
| 15 | 559.25 | 11.5 | 0.5 | 38 | 898.25 | 24.5 | 0.5 |
| 16 | 570.25 | 10.5 | 0.5 | 39 | 922.25 | 26.5 | 0.5 |
| 17 | 580.25 | 11.5 | 0.5 | 40 | 948.25 | 28.5 | 0.5 |
| 18 | 591.25 | 12.5 | 0.5 | 41 | 976.25 | 31.5 | 0.5 |
| 19 | 603.25 | 12.5 | 0.5 | 42 | 1007.25 | 35.5 | 0.5 |
| 20 | 615.25 | 12.5 | 0.5 | 43 | 1042.25 | 41.5 | 0.5 |
| 21 | 627.25 | 11.5 | 0.5 | 44 | 1083.25 | 50.5 | 0.525 |
| 22 | 638.25 | 13.5 | 0.5 | 45 | 1133.225 | 516.8 |  |
| 23 | 651.25 | 12.5 | 0.5 |  |  |  |  |

### Instrument parameters

**Table 2:** Instrument parameters.

| MS survey scan |  |
| --- | --- |
| Spray Source | Positive mode |
| Spray voltage | 2.650kV |
| Ion transfer tube Temperature | 250°C |
| MS1 Orbitrap resolution | 120000 |
| MS2 Orbitrap resolution | 30000 |
| Full MS Scan Cycle | 350-1650 m/z |
| RF Lens (%) | 40 |
| AGC (%) | 300 |
| maximum injection time mode | Custom |
| Maximum injection time | 20 ms |
| Source fragmentation | disabled |
| **MS survey scan followed by MS/MS DIA Scan events** |  |
| Multiplex ions | False |
| Collision energy mode | Stepped |
| Collision energy type | Normalized |
| HCD collision energies (%) | 25.5,27,30 |
| Orbitrap resolution | 30000 |
| First mass | 200 |
| RF lens (%) | 40 |
| AGC target | Custom |
| Normalized AGC target (%) | 3000 |
| Maximum injection time mode | Custom |
| Maximum injection time | 55 ms |

### DIA data processing

DIA mass spectrometry data was processed in Spectronaut 16 and 17. The default search parameters, along with optimised more stringent parameters were used. The tables bellow specify the identification setting details.

**Table 3:** Default Spectronaut Identification Settings.

| **Default Identification Settings** |  |
| --- | --- |
| Precursor Q-value Cutoff | 0.01 |
| Precursor PEP Cutoff | 0.2 |
| Protein FDR Strategy | Accurate |
| Protein Q-value Cutoff (Experiment) | 0.01 |
| Protein Q-value Cutoff (Run) | 0.05 |
| Protein PEP Cutoff | 0.75 |

**Table 4.** Stringent Spectronaut Identification Settings.

| **Optimized Identification Settings** |  |
| --- | --- |
| Precursor Q-value Cutoff | 0.01 |
| Precursor PEP Cutoff | 0.01 |
| Protein FDR Strategy | Accurate |
| Protein Q-value Cutoff (Experiment) | 0.01 |
| Protein Q-value Cutoff (Run) | 0.01 |
| Protein PEP Cutoff | 0.01 |

### Data normalisation

The median intensities were calculated for each sample after filtering out any proteins with an intensity of 0. The individual protein intensities across each sample were divided by the sample median. Volcano plots were made using -Log_10_ (P-value), and Log_2_ Fold Change (spiked/non-spiked) samples. P-values and fold changes values were calculated using the bioconductor package, Limma (32), after Log_2_ transforming the normalised intensity data. The p-value cut-offs are set to 0.01, on denoted as a horizontal grey line, at 2 on -Log_10_ scale, and the Fold-change cut-off was set to 2, and −2, denoted as 1 and −1 on the Log_2_ scale on the volcano plot.

### Theoretical Tryptic Digest

Protein sequences were downloaded from UniProt in January, 2023. *Mus musculus* C57BL/J6 sequences including isoforms and variant sequences were obtained by filtering for ‘Mouse’ in the ‘model_organism’ field and selecting only reviewed records, and the ‘canonical and isoforms’ download option. *C. albicans* SC5314 sequences were selected using the NCBI taxonomy identifier 237561, and again both canonical and isoform sequences downloaded. In-silico trypsin digests were carried out using the ‘pepdigest’ tool from the EMBOSS 6.6.0.0 package (33), assuming complete digestion and cutting only at favoured sites (K/R not followed by K,R,I,F,L or P) and a peptide length >6 amino acids.

The resulting peptides were parsed using custom Python code and pandas 1.5.3 (34) data frames constructed for both organisms. Four categories of peptide were identified using combinations of pandas concat(), unique() and drop_duplicates() methods to create subsets of peptides:

1) Unique within each organism i.e. only occurring once within Mouse or *C. albicans*
2) Shared within each organism, i.e. peptides that map to multiple proteins within the same organism
3) Unique to each organism
4) Shared between the two organisms (referred to as cross-species peptides), i.e. peptides that map to proteins from both Mouse and *C. albicans*

### Code availability

All code was produced in Jupyter notebooks and is freely available under an MIT license from https://github.com/bartongroup/Mouse-C.albicans-Peptide-Overlaps, and interactively through MyBinder at https://mybinder.org/v2/gh/bartongroup/Mouse-C.albicans-Peptide-Overlaps/HEAD.

### FASTA files

A *Mus musculus* SwissProt canonical with isoforms (February 2022) database and *C. albicans* TrEMBL (May 2023) database were used for the Spectronaut searches.

## Results

### Mouse and *C. albicans* proteomes contain cross-species peptides

To understand the theoretical homology between the host *Mus musculus* (Mouse) proteome and pathogenic *Candida albicans (C. albicans)* proteome, we performed an *in-silico* tryptic digest for both species (see methods). A comparison of their protein databases revealed that the mouse FASTA contained a total of 25,489 proteins, while the *C. albicans* FASTA contained 6,042 proteins (Fig. 1A). An *in-silico* tryptic digest, which excluded peptides with less than 6 amino acids, resulted in 451,723 theoretical Mouse peptides of which 144,713 peptides are shared between more than one Mouse protein and 132,462 *C. albicans* peptide sequences, with 1,261 between more than one *C. albicans* protein (Figure 1B). We next sought to evaluate the number of peptides that were shared between Mouse and *C. albicans*, referred to as cross-species peptides, to determine the potential for misidentifications. In total there were only 351 cross-species peptides shared between Mouse and *C. albicans.* This meant that only 0.08% of all Mouse peptide sequences were shared with *C. albicans*, and 0.26% of all *C. albicans* peptides were shared with Mouse (Figure 1C).

**Figure 1.**
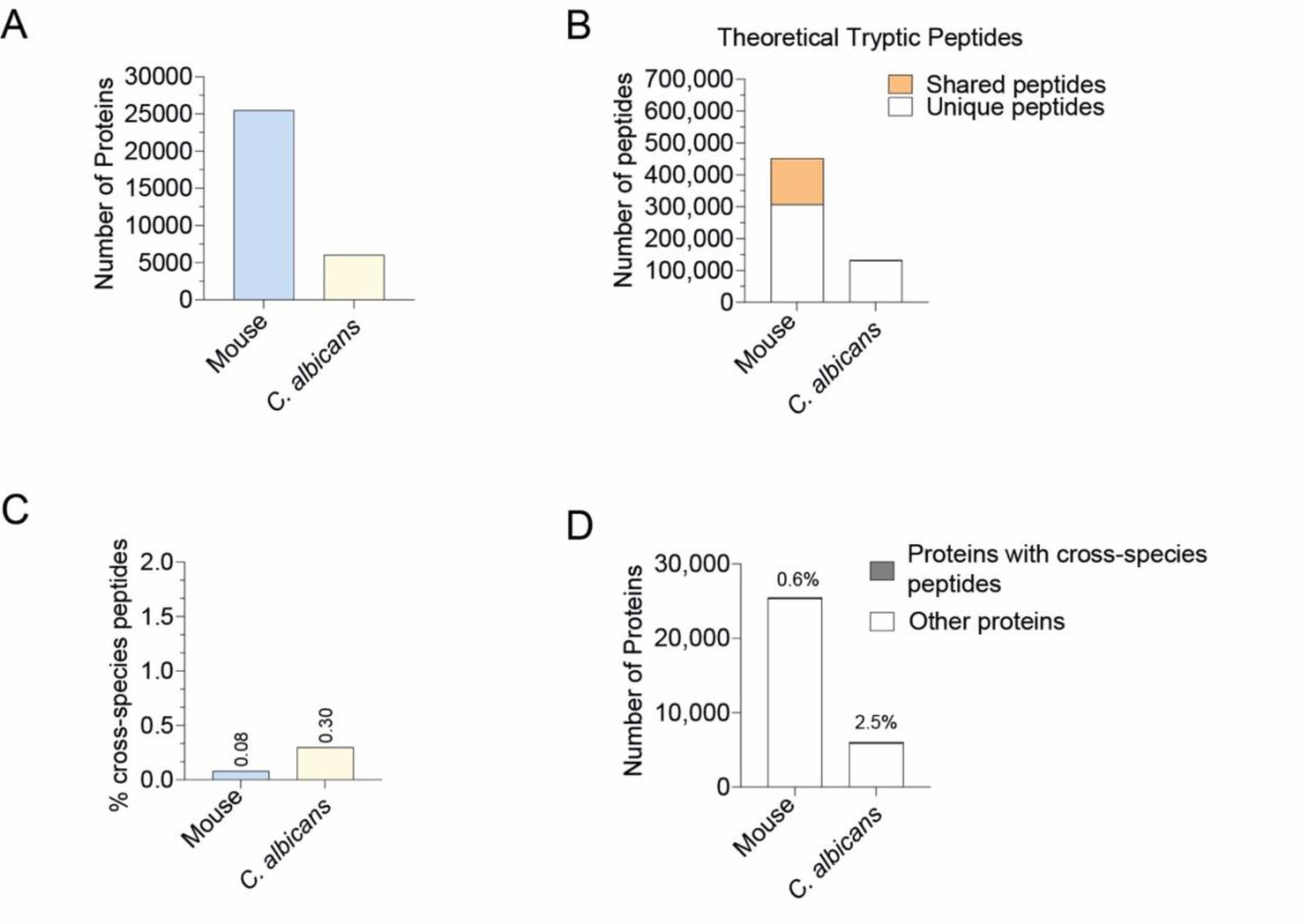
Homology between Mouse and C. *albicans*: **(A)** Number of proteins present in the FASTA file entries for Mouse and *C. albicans.* **(B)** Total number theoretical tryptic peptide sequences found in Mouse and *C. albicans* species. Shared peptides are highlighted in orange. Only peptides with 7 amino acids or more were considered. **(C)** The percentage of cross-species peptides present in the Mouse and *C. albicans* peptidome. **(D)** Number of proteins in Mouse and *C. albicans.* Proteins containing cross-species peptides highlighted in grey.

These cross-species peptides mapped to 148 Mouse proteins, representing 0.6% of the entire Mouse proteome (Figure 1D). As a result, when quantifying the Mouse proteome via mass spectrometry in *C. albicans* infected samples there is limited potential for miss-assignment of *C. albicans* peptides as Mouse peptides. This implies that only a small subset of C. albicans peptides could be wrongfully identified as Mouse peptides and assigned to Mouse proteins, hence adding quantitative noise.

### Including a FASTA file for both species reduces quantitative disruptions

To experimentally address if the presence of *C. albicans* peptides affects the quantification of Mouse proteins, we generated lysates from cultures of Mouse Bone Marrow Derived Macrophages (BMDMs) and separately *C. albicans* lysates. Three biological replicates of BMDMs, were separated into two aliquots. One of the aliquots was spiked after lysis with *C. albicans* at a 1:4 ratio, referred to as ‘spiked’, the other was left without any spike-ins, referred to as ‘Non-spiked’. We then used a DIA-based analysis to characterise the proteome of the Mouse BMDMs, comparing spiked and non-spiked samples (shown in Figure 2A), with the raw files being processed with Spectronaut (8).

**Figure 2.**
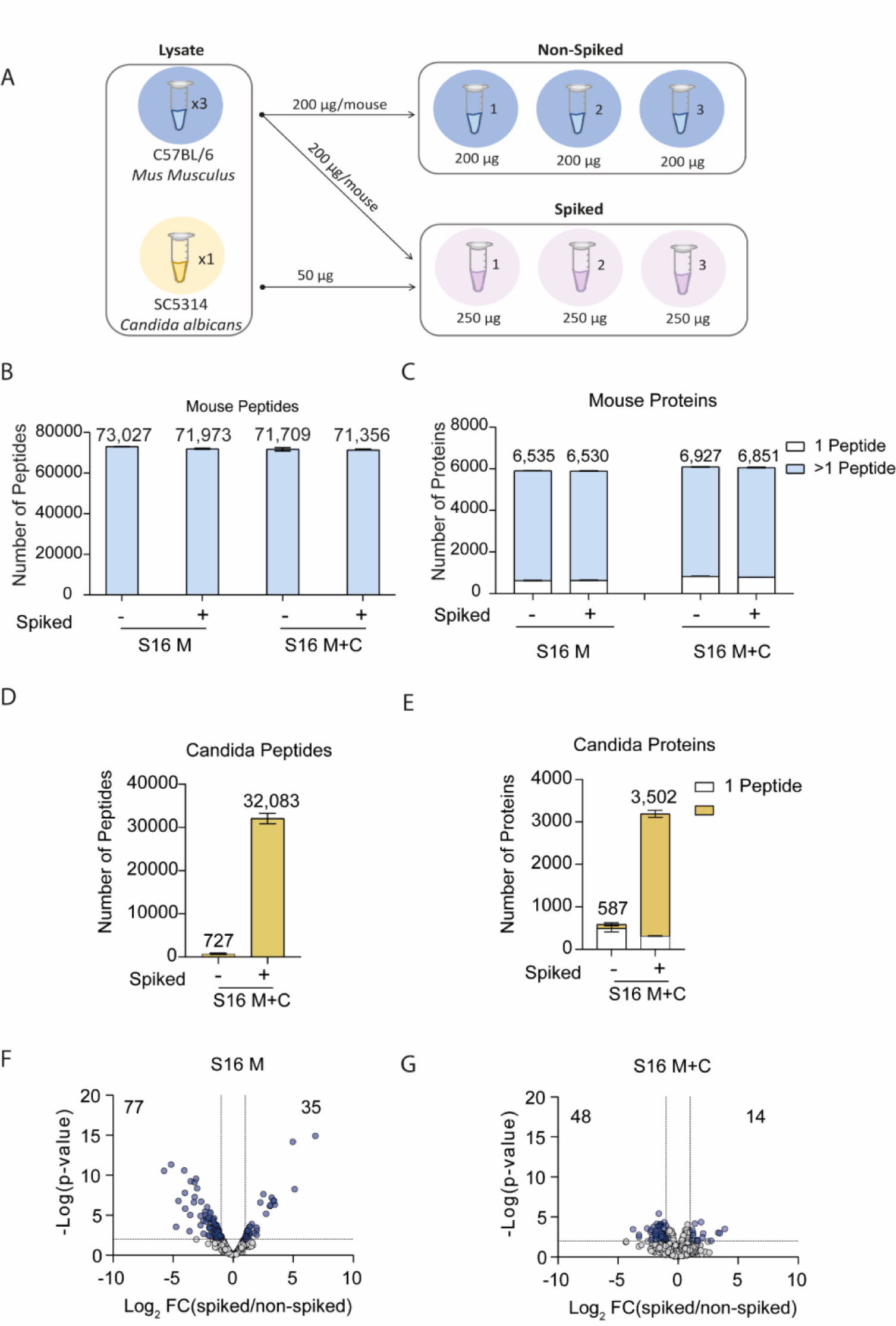
Identification and quantification errors: **(A)** Schematic of experiment where Mouse BMDMs and *C. albicans* were lysed separately for proteomics, and subsequently *C. albicans* lysates were spiked with the BMDM lysates. (Mouse n=3, *C. albicans* n=1). Number of **(B)** Mouse peptides, **(C)** Mouse proteins, identified using either a Mouse only (M) or Mouse and *C. albicans* (M+C) FASTA. Proteins identified from only one peptide are highlighted in white. Number of **(D)** *C. albicans* peptides and **(E)** C. albicans proteins identified using the Mouse and *C. albicans* (M+C) FASTA. Volcano plots showing the Log_2_ fold change of the spiked versus the non-spiked samples and the-Log_10_(p-value) for **(F)** Spectronaut 16 (S16) using default identification settings and a Mouse only FASTA, **(G)** Spectronaut 16 (S16) using default identification settings and a Mouse and a *C. albicans* FASTA. Proteins with a p-value<0.01 and a fold change >2 are highlighted in blue.

We first set out to determine if it was beneficial to include both a FASTA file for Mouse and a FASTA file for *C. albicans* within the search space. We found no major differences in the number of Mouse peptides identified when using only a Mouse FASTA or using both a Mouse and a *C. albicans* FASTA file. (Fig. 2B). This also proved true when analysing the number of mouse proteins identified (Fig. 2C). We also checked the number of *C. albicans* peptides identified and unsurprisingly, none were present if the FASTA is not included. We noticed however that when we included the *C. albicans* FASTA, we detected *C. albicans* peptides (Figure 2 D) and proteins (Figure 2 E) not only in the spiked but also the non-spiked samples, where they should not have been present.

We next set out to determine the quantitative differences of using only a Mouse FASTA or both a Mouse FASTA and a C. albicans FASTA. For this we performed a differential expression analysis on the mouse proteome comparing the spiked and non-spiked samples. It is important to highlight the *C. albicans* lysate was spiked after the BMDM samples were lysed, hence no biological differences are expected between the spiked and non-spiked samples, though some technical differences can be present. The data showed that when applying a fold change > 2 and p-value < 0.01 cut-off, 112 proteins were changed between the spiked and non-spiked BMDMs when using only a Mouse FASTA (Figure 2F). We inferred this scenario was potentially caused by the assignment of the intensity of *C. albicans* peptides into Mouse proteins. Interestingly, only 2 of these proteins contained a cross-species peptide between Mouse and *C. albicans*, suggesting that there may also be an issue with incorrect peptide assignments. We hypothesised that also including the *C. albicans* FASTA file had the potential to improve the quantitative data and minimise the assignment of intensity from *C. albicans* peptides into Mouse proteins. The addition of the *C. albicans* FASTA reduced the number of significantly changed proteins to 62 (Figure 2G), a reduction of 45%. We determined that including the *C. albicans* FASTA file was important to minimise the quantification errors. Hence for all the subsequent analyses a Mouse and a *C. albicans* FASTA were used.

### Optimising Spectronaut identification settings minimises misidentifications while maintaining proteome depth

Having determined it was optimal to use both the Mouse and *C. albicans* FASTA files (M+C) we focussed on the potential false positives and misidentifications seen in the non-spiked samples (Figure 2D-E). For this work we used Spectronaut (S) version 16 and 17 (S16 or S17, respectively) (schematic of analysis in Figure 3A). We first compared both version 16 and 17 using the default settings. In this context we focussed our analysis on the number of *C. albicans* peptides and proteins that were identified both within the *C. albicans* spiked samples and the non-spiked samples. We first focussed on the spiked samples. Here, S16 identified a total of 32,083 *C. albicans* peptides that matched to 3,190 total proteins identified (Figure 3B-C). Those same samples analysed with S17 using the same default settings, identified 39,413 peptides, which represented a 20.5% increase, and these peptides matched to 3,477 proteins, which represented 8.6% increase in the number of proteins identified. This suggests that S17 was the superior option with respect to peptide and protein identification rates.

**Figure 3.**
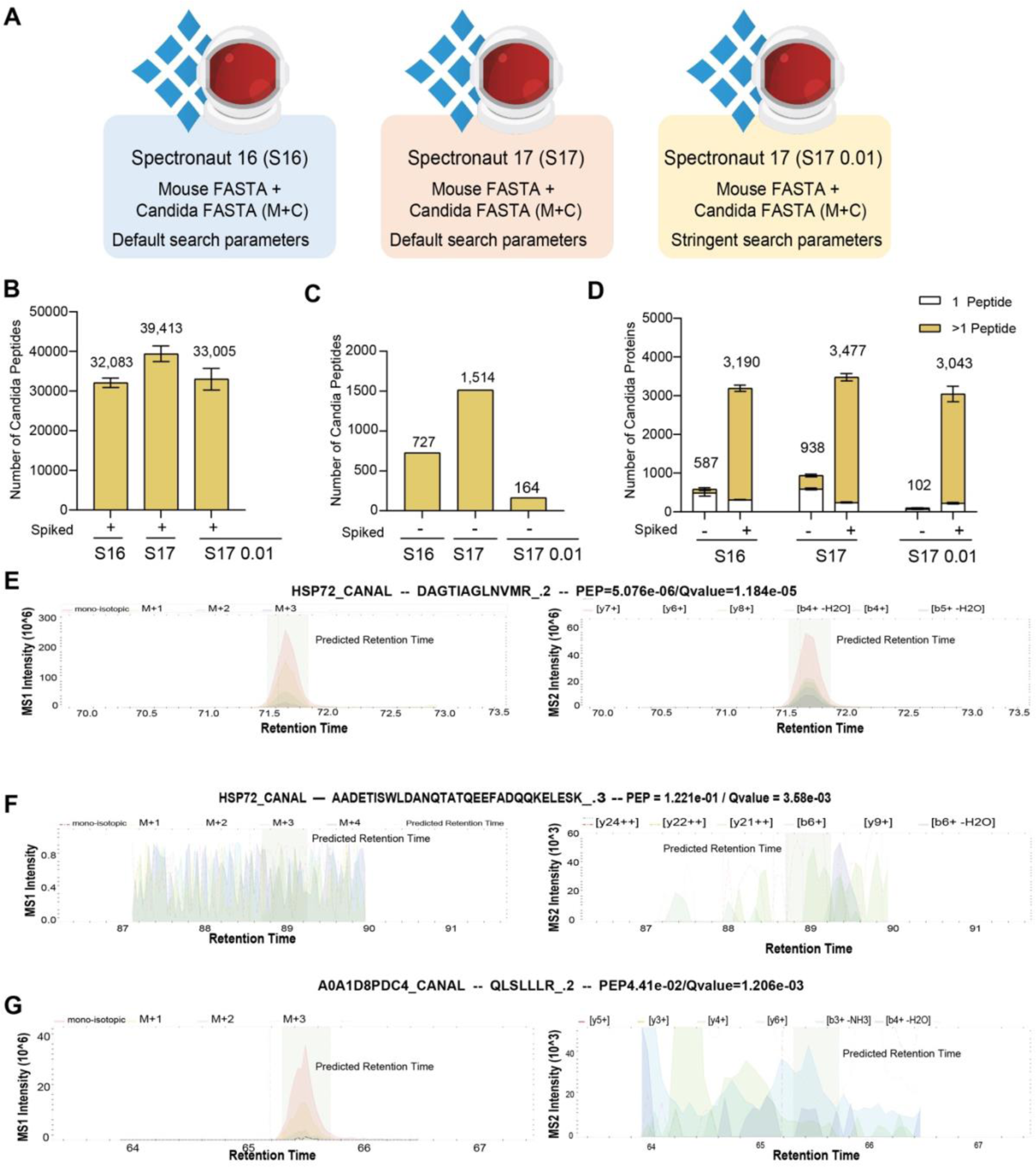
*C. albicans* peptide and protein identification across Spectronaut versions and settings: (**A)** Schematic of the data processing pipelines generated using Spectronaut 16 (S16) or 17 (S17), with default or stringent settings. The number of *C. albicans* peptides detected across the different pipelines in the **(B)** spiked samples **(C)** non-spiked samples. (D) Number of *C. albicans* proteins detected in both spiked and non-spiked samples. Proteins identified with a single peptide are coloured in white. Peptide (MS1) and precursor ion (MS2) eXtracted Ion Chromatogram (XIC) profiles for (E) protein HSP72_CANAL with precursor DAGTIAGLNMR_.2, (F) protein HSP72_CANAL with precursor AADETISWLDANQTATQEEFADQQKELESK_.3, and (H) protein A0A1D8PDC4_CANAL with precursor QLSLLLR_.2.

We next focussed on *C. albicans* proteins and peptides that were detected in the non-spiked samples where only Mouse lysate was present. *C. albicans* proteins and peptides that were identified in the non-spiked samples provided examples of potential false positives. Here, we found that S16 identified 727 *C. albicans* peptides in the non-spiked samples, mapping to 587 proteins. S17 again displayed higher identification rates, with a mean of 1,514 *C. albicans* peptides matched to 938 proteins in the non-spiked samples. Across all 3 replicates a total of 1,538 *C. albicans* proteins were identified in at least 1/3 replicates. We found only 94 out of the 1538 identified by S17 were cross-species peptides, suggesting the majority of these peptides should not be present in the non-spiked samples and thus are considered to be false positives. This scenario is particularly problematic as the false positives at the protein level represent ∼16% of the *C. albicans* proteome, which can lead to inaccurate biological interpretations.

We suspected the previously seen false positives were caused by the erroneous transfer of identifications across the different runs, a similar situation is seen in DDA with ‘match between runs’ (36, 37). Hence, we decided to optimise the default search settings in Spectronaut to minimise the number of false positives. We increased the stringency of the default settings by setting the posterior error probability to ≤ 0.01 and by setting the protein q-value scores across the runs to ≤ 0.01 (analysis settings denoted S17 M+C 0.01).

Our data showed that the updated parameters were extremely effective at removing false identifications in the non-spiked samples, reducing the mean number of *C. albicans* peptides detected from 1,514 to 164, and the mean number of *C. albicans* proteins from 938 to 102.

Across all 3 non-spiked samples, a total of 164 *C. albicans* proteins were detected; 89 of which contained cross-species peptides. This suggested that for 54% of these proteins, the issues were not caused by spurious peptide identifications, but incorrect peptide to protein group assignment. The data also revealed that the majority of *C. albicans* peptides that were still detected with the optimised parameters had clear extracted ion chromatogram (XIC) peaks at the predicted retention times (Figure 3E and supplementary Figure 1-2). Whereas the identifications that were filtered out frequently displayed no concrete precursor ion XCI peaks (Figure 3F-G and supplementary Figure 3-4 for each replicate spectra).

**Figure 4.**
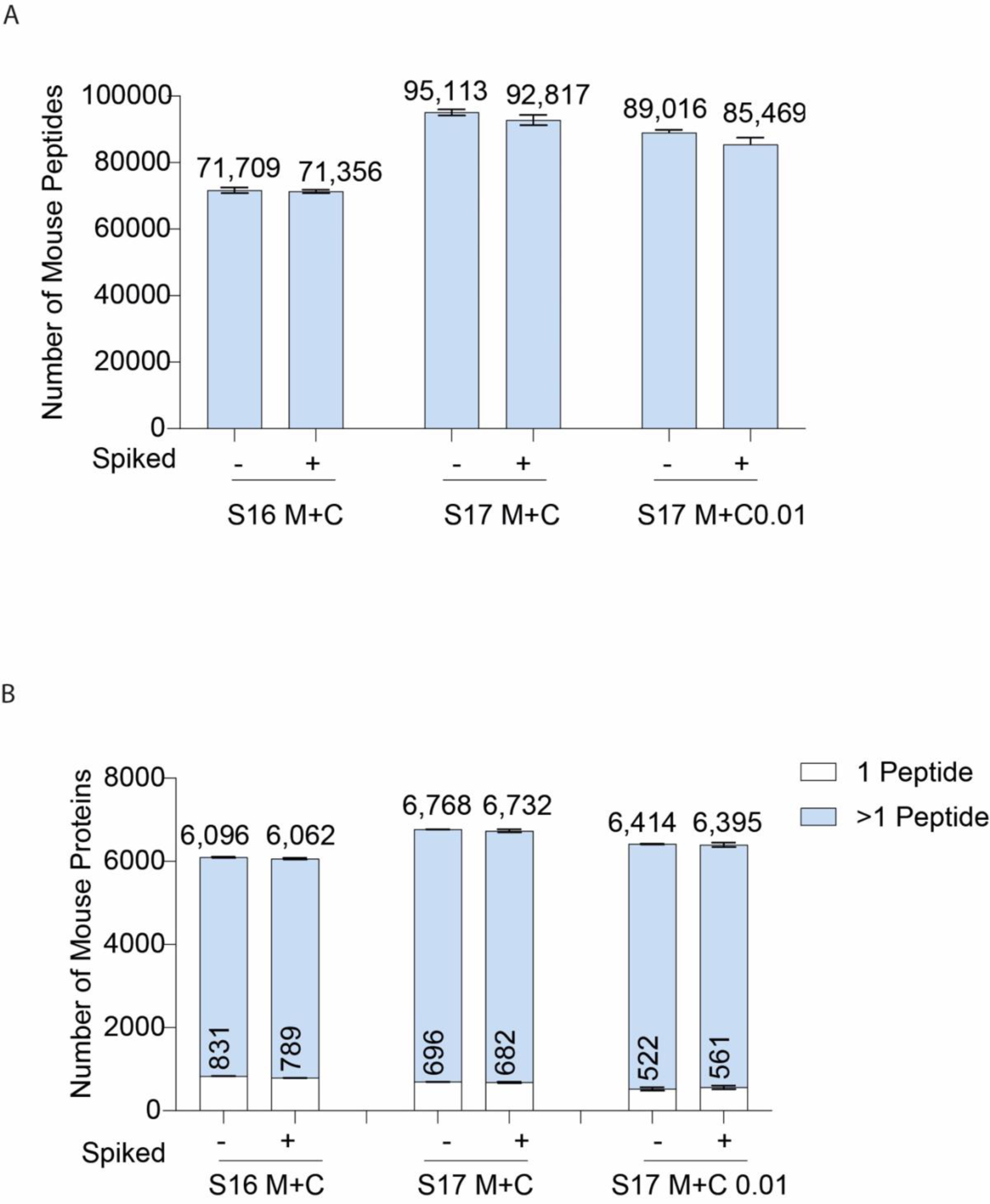
Mouse identifications across Spectronaut settings: Number of **(A)** Mouse peptides and **(B)** Mouse proteins; identified in spiked and non-spiked samples. Proteins identified with one peptide are noted in white.

We also wanted to understand the effect that the stringent parameters would have on Mouse proteome coverage, i.e. Mouse peptides found in the non-spiked samples, as we suspected the stringent parameters might cause a considerable reduction in the number of these peptides. The standard setting on S17 led to the identification of 95,113 Mouse peptides and 6,768 proteins in the non-spiked samples (Figure 4 A and B). The use of the stringent settings only led to a reduction of 6.6% in the number of mouse peptides identified (89,016 peptides) in this sample which translated into a 5.4 % reduction in the number of Mouse proteins (6,414 proteins) identified. Furthermore, the number of Mouse proteins identified with the stringent parameters in S17 was still higher than the standard parameters on S16, suggesting these optimised setting on the newer versions of Spectronaut are a very effective strategy to obtain increased data quality at a small penalty. Additionally, the quantification of Mouse proteins highlighted as significantly changed due to the presence of *C. albicans*, of which just a small proportion had cross-species peptides, was reduced by >50% in S17 0.01, while still being detected within the dataset (Supplementary Figure 5).

Overall, although we did not entirely remove misidentifications with our optimised settings, we were able to make significant improvements without a considerable reduction of proteome depth.

## Discussion

DIA-based proteomic workflows are characterised by having a much higher complexity spectra than what is usually encountered in the narrow DDA isolation window (3–5). The computational solutions to analyse such complex data originally required specific peptide or spectral libraries to be generated (6–8). These libraries were generated in DDA and added cost and complexity to DIA projects. A big breakthrough towards the widespread implementation of DIA occurred with the implementation of library free searches across the different software tools (38–40). This meant it was no longer necessary to generate expensive DDA-based libraries. However, the complexity of the spectra is still present, and though library free search algorithms leverage deep neural networks to improve the matching and deconvolution (41, 42), some issues remain.

In immunological research, it is not uncommon to study infection models, where a host, i.e. *Mus musculus*, is infected by a pathogen such as *C. albicans* (20, 21). Hence, we became interested in understanding the potential complications that could arise when analysing these two proteomes within a single sample. This scenario represents two heterogenous populations where some proteins should only be present in one of the two populations, which frequently occurs in proteomic studies. Here we compared the proteomes of Mouse macrophages that were either spiked with *C. albicans* post lysis or not spiked.

Mouse and *C. albicans* displayed a low level of sequence homology, which suggested limited issues relating to peptide misidentifications across the two species. However, our data revealed that using the default search settings 1,514 peptides and 938 proteins were identified in the non-spiked murine samples, representing ∼16% of the *C. albicans* proteome. All the 1,514 peptides identified in the non-spiked samples were detected in *C. albicans* spiked samples, and the majority had robust PEPs and XIC peaks. This suggests that the peptides were correctly identified in those samples. The issue with the C. albicans peptides detected in the non-spiked samples centred around the transfer of these identifications across the different runs; an issue that has been described in DDA with ‘match between runs’ algorithms (36, 37, 43).

Thus, we optimised the search parameters to increase the stringency. Firstly, we set the protein Q-value cut-off per run to <0.01, this would increase the strictness of the identification transfers across the different samples. Secondly, we set the posterior error probability (PEP) at the precursor and protein level to <0.01, increasing the stringency of the overall identifications. These updated parameters proved extremely successful at improving the quality of the identifications, reducing the number of false positive identifications of *C. albicans* peptides and proteins present in the non-spike samples, by ∼90%. Importantly, this increased stringency came at a limited cost; where the total mouse peptide identifications, which we will call true positives, were only reduced by 6.4%. We believe our work is not limited to multispecies but would also be of great value when analysing heterogenous populations within the same experiment. For example, a DIA study looking at multiple distinct cell types or studying different tissues would also greatly benefit from the previously mentioned reduction in false positives.

As newer instruments and analytical methods enable a more comprehensive coverage while using shorter gradients (1, 2, 9), we believe the focus should migrate from maximising the number of identifications, to increasing the quality of such identifications. We show that a way to achieve this when using Spectronaut only requires modifying a small subset of search parameters, which results in dramatically reduced false positives with only a minor reduction in overall proteome depth.

It is also worth noting that our work manually searched a subset of peptides and fragment ions which we knew should be absent from specific populations. This search discovered multiple matches without discernible XIC peaks. It would also be of great benefit to develop a tool that systematically analysed the peptide and precursor identifications, flagging up the number of peptides and precursor ions without discernible XIC peaks within the expected retention times. If this gets integrated into an automated reporting tool, it will enable a much easier and user-friendly approach to monitor DIA data quality.

## Acknowledgements

We would like to thank Doreen Cantrell for her support, Andrew Howden and Tony Li for their insights and helpful discussions, all the members of the Cantrell and the Arthur group, the University of Dundee Biological Resources Unit, Flow Cytometry Facility (R. Clarke and team) and Fingerprints Proteomics Facility (D. Lamond and team). This work was funded by a Wellcome Trust PhD studentship to Christa Baker (102132/B/13/Z/WT), a Wellcome Trust Principal Research Fellowship (205023/Z/16/Z) and Strategic Award (105024/Z/14/Z) to Doreen Cantrell.

## Data availability

All raw files, Spectronaut sne files, Spectronaut reports, FASTA files and the experimental template have been uploaded to PRIDE(35) under accession number PXD045958 (https://www.ebi.ac.uk/pride/archive/projects/PXD045958).

## Author Contributions

AJB, CPB and JSCA conceived the project. CPB performed the macrophage and *C. albicans* culture and sample processing. AJB performed all Spectronaut searches. JA performed the theoretical peptide analysis. AJB, CPB and RB analysed the data. CPB generated all figures, AJB edited them. AJB and CPB wrote the manuscript with input from all authors. AJB and JSCA supervised the project.

## Supplementary data

**Supplementary Figure 1:**
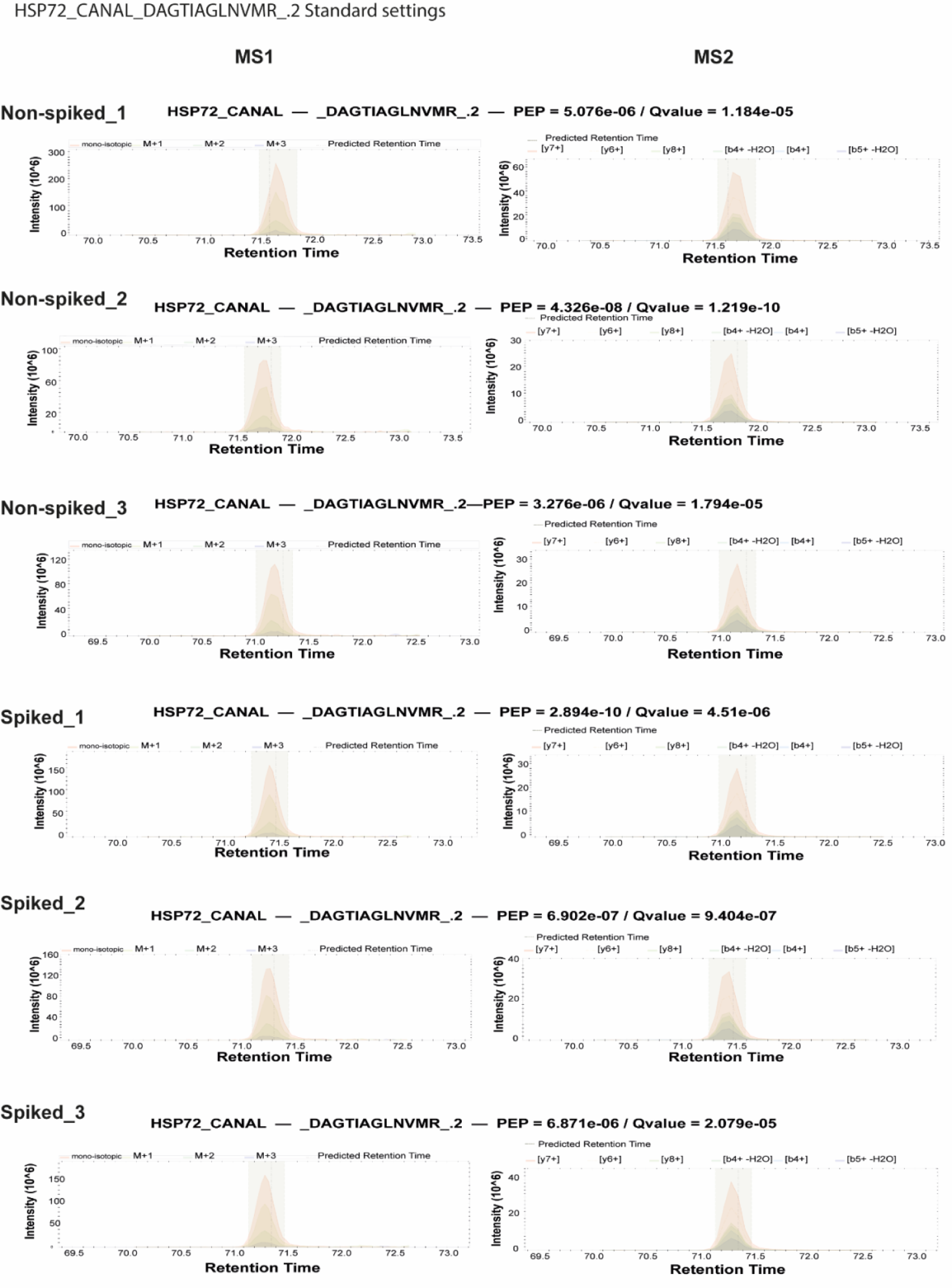
Spectronaut 17 default settings MSI and MS2 extracted ion chromatogram (XIC) profiles for *C. albicans* proteome protein HSP72 and precursor DAGTIAGLNVMR_.2, across all samples, both spiked and non-spiked.

**Supplementary Figure 2:**
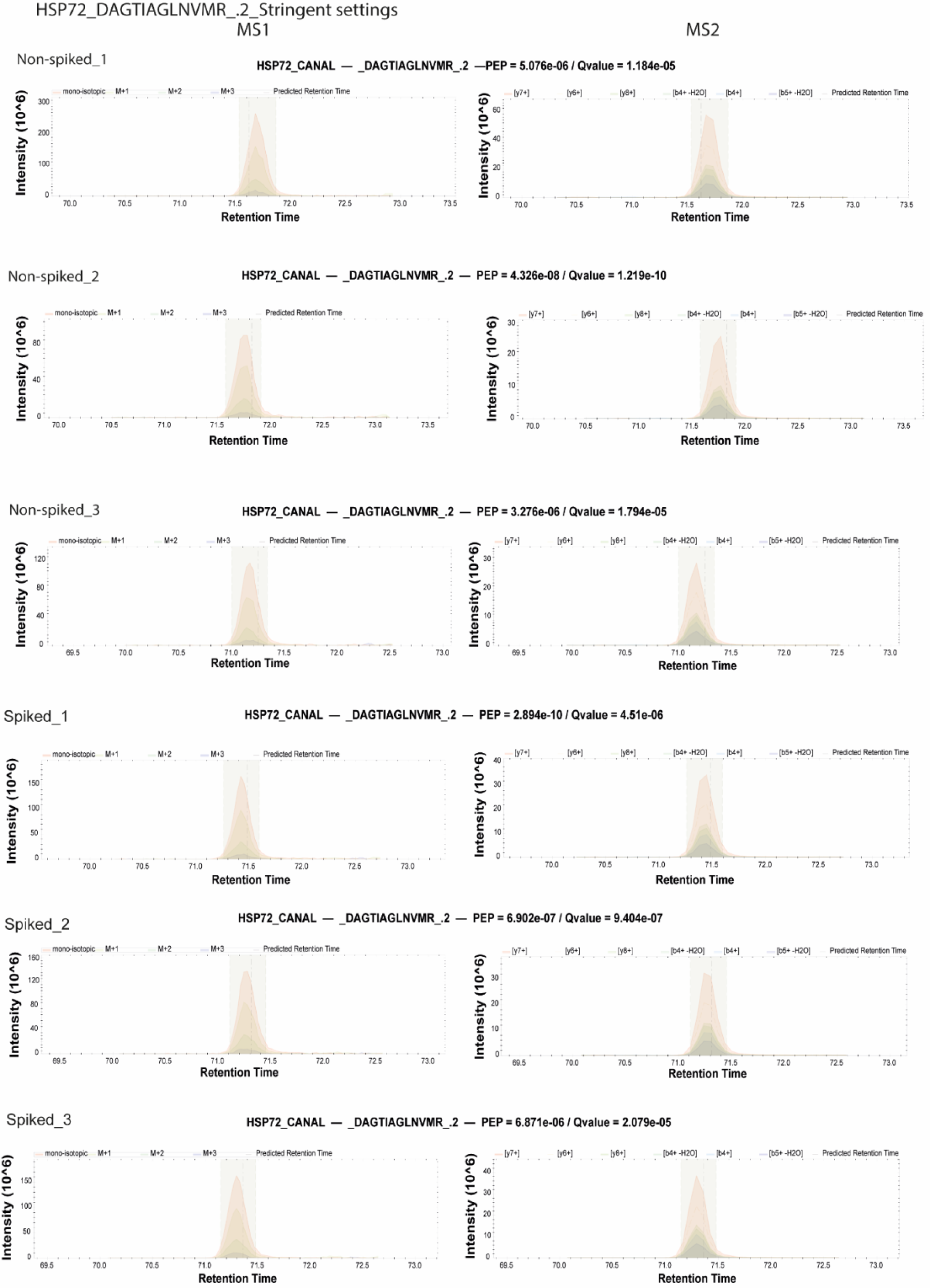
Spectronaut 17 stringent settings MSI and MS2 extracted ion chromatogram (XIC) profiles for *C. albicans* proteome protein HSP72 and precursor DAGTIAGLNVMR_.2, across all samples, both spiked and non-spiked.

**Supplementary Figure 3:**
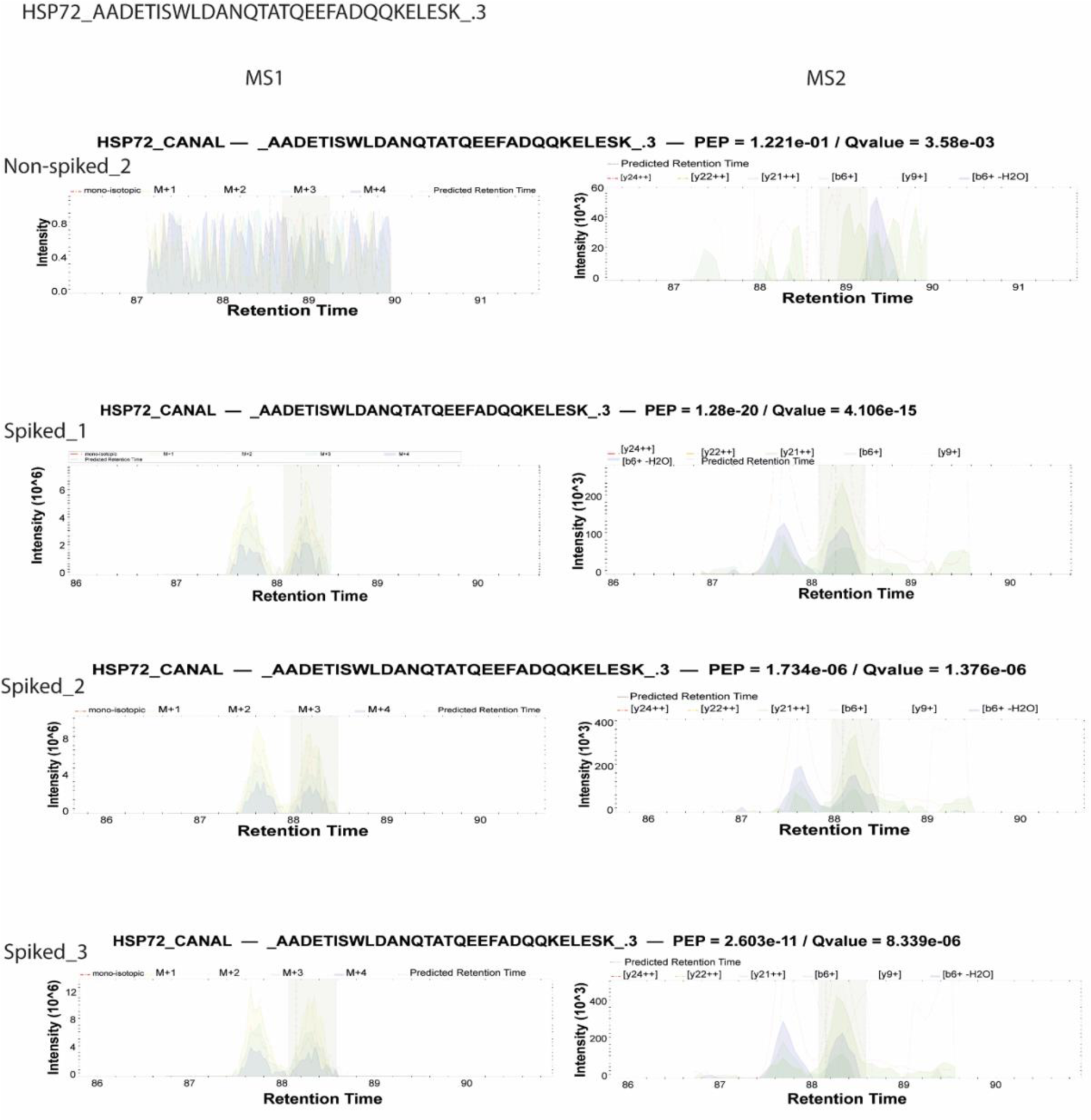
Spectronaut 17 default settings MSI and MS2 extracted ion chromatogram (XIC) profiles for *C. albicans* proteome protein HSP72 and precursor AADETISWLDANQTATQEEFADQQKELESK_.3, across all samples, both spiked and non-spiked.

**Supplementary Figure 4:**
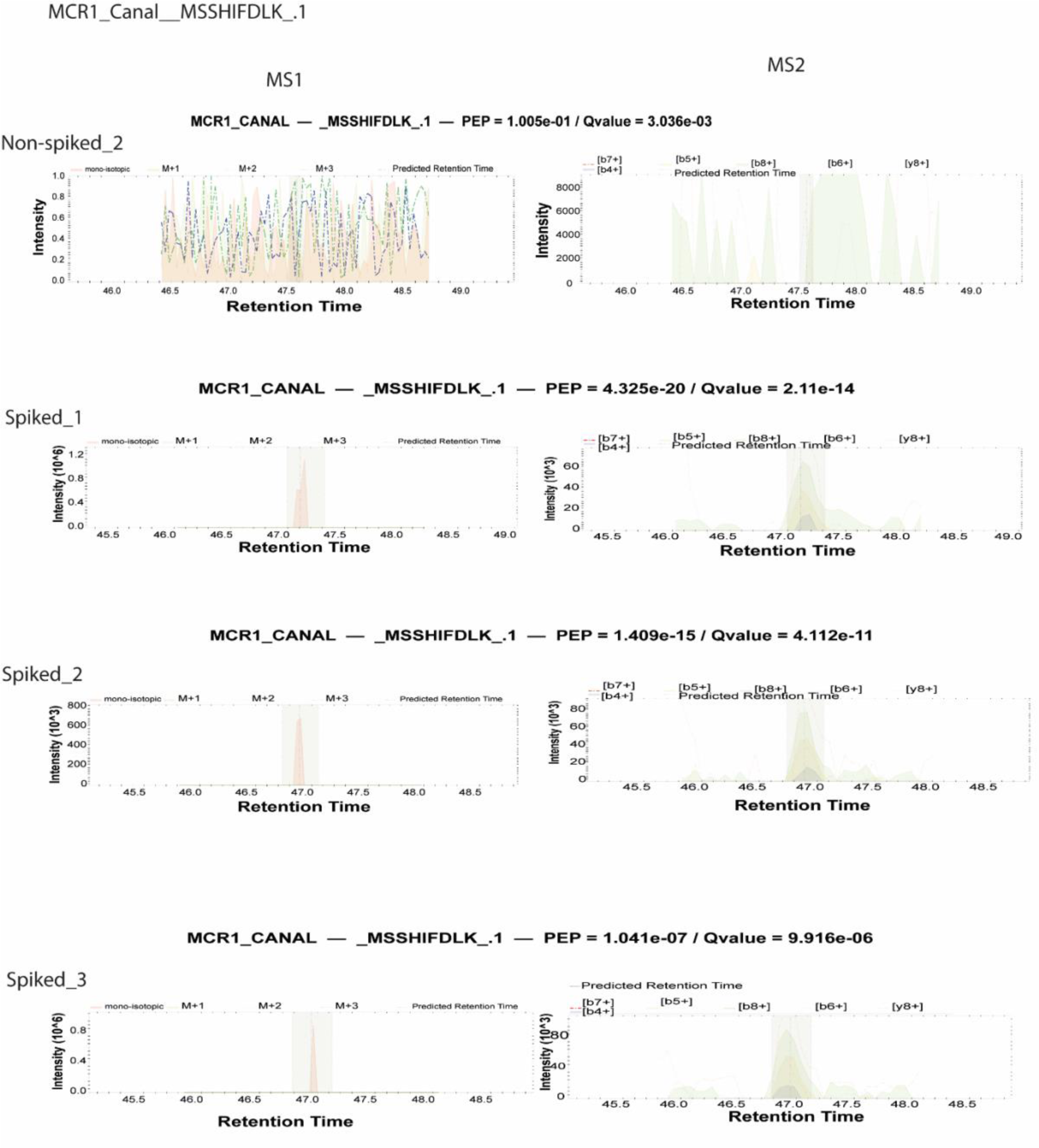
Spectronaut 17 default settings MSI and MS2 extracted ion chromatogram (XIC) profiles for *C. albicans* proteome protein HSP72 and precursor MSSHIFDLK_.1, across all samples, both spiked and non-spiked.

**Supplementary Figure 5.**
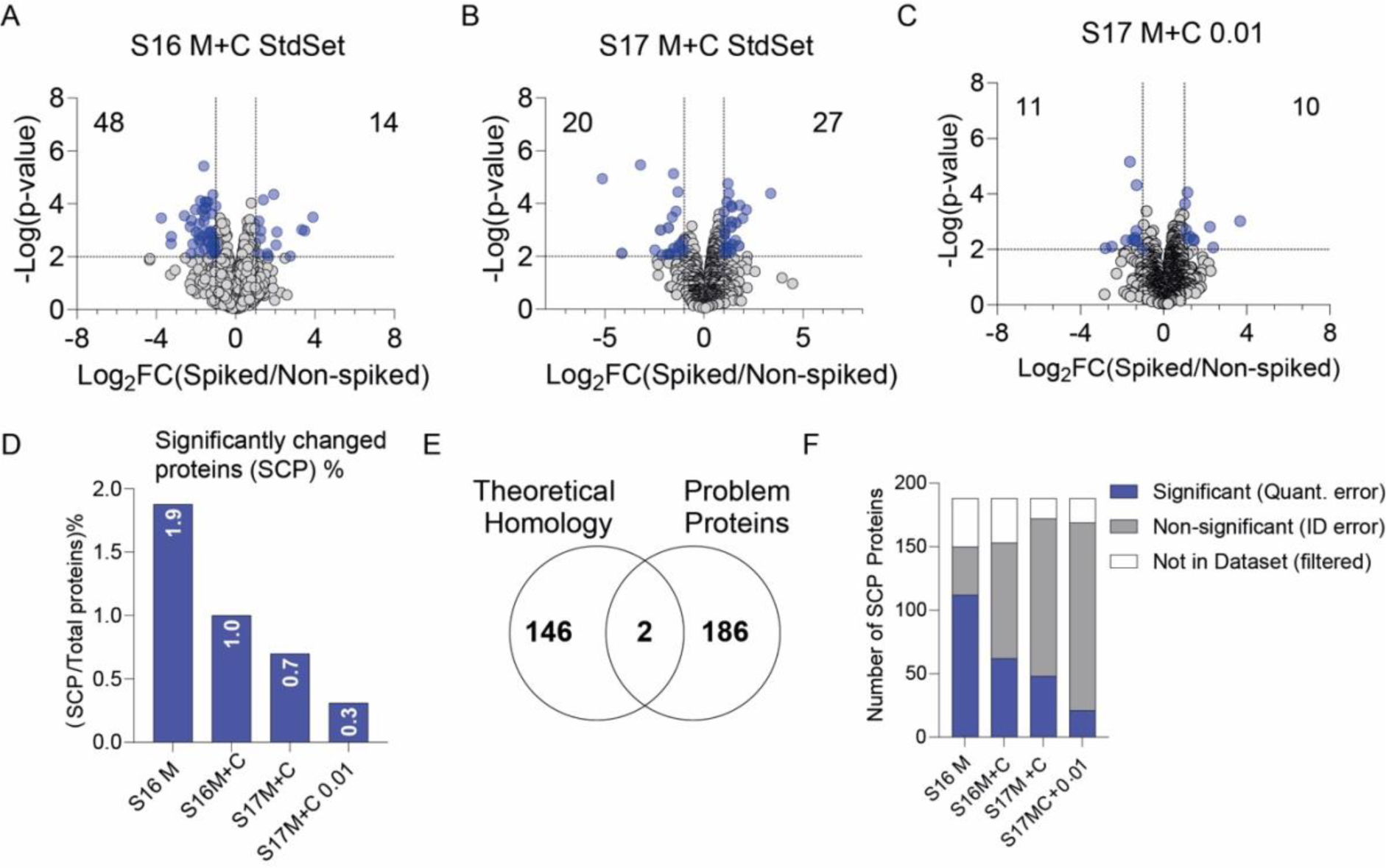
Volcano plots show the Log_2_ fold change (spiked vs non-spiked), and the -Log_10_ p-value and highlights proteins with a p-value< 0.01 and −2>FC>2. All volcano plots Spectronaut analysis includes Mouse and *C. albicans* FASTA files. **(A)** Spectronaut 16, default settings, **(B)** Spectronaut 17, default settings and **(C)** Spectronaut 17 with 0.01 identification settings. (D-E) The significant proteins from each analysis were pooled to make a compiled list of “problematic proteins” p-value <0.01 and −2>FC>2. **(D)** Plots problematic proteins in each analysis as a percentage of the total proteins identified in the dataset. **(E)** compares this list of Mouse problematic proteins against the theoretical homology between Mouse and *C. albicans* to determine how many of these errors are due to homology or misidentification while **(F)** plots the problematic proteins, highlighting the number of significant or non-significant proteins for that analysis, as well as proteins that were not identified in the analysis.

